# Regulation of stem cell identity by miR-200a during spinal cord regeneration

**DOI:** 10.1101/2021.07.21.453081

**Authors:** Sarah E. Walker, Keith Z. Sabin, Micah D. Gearhart, Kenta Yamamoto, Karen Echeverri

## Abstract

Axolotls are an important model organism for multiple types of regeneration, including functional spinal cord regeneration. Remarkably, axolotls can repair their spinal cord after a small lesion injury and can also regenerate their entire tail following amputation. Several classical signaling pathways that are used during development are reactivated during regeneration, but how this is regulated remains a mystery. We have previously identified miR-200a as a key factor that promotes successful spinal cord regeneration. Here, using RNA-seq analysis, we discovered that the inhibition of miR-200a results in an upregulation of the classical mesodermal marker *brachyury* in spinal cord cells after injury. However, these cells still express the neural stem cell marker *sox2*. *In vivo* lineage tracing allowed us to determine that these cells can give rise to cells of both the neural and mesoderm lineage. Additionally, we found that miR-200a can directly regulate *brachyury* via a seed sequence in the 3’UTR of the gene. Our data indicate that miR-200a represses mesodermal cell fate after a small lesion injury in the spinal cord when only glial cells and neurons need to be replaced.

**Summary Statement:** After spinal cord injury, miR-200 fine-tunes expression levels *brachyury* and *β-catenin* to direct spinal cord stem into cells of the mesodermal or ectodermal lineage.

## Introduction

Regeneration has been observed throughout the plant and animal kingdoms for many years (Sanchez Alvarado and Tsonis, 2006). Among vertebrates, the Mexican axolotl salamander has the remarkable ability to faithfully regenerate its spinal cord after injury. This process has been most commonly studied in the context of tail amputation (McHedlishvili et al., 2012, Monaghan et al., 2007, Piatt, 1955, Rodrigo Albors et al., 2015, Zhang et al., 2000, Zhang et al., 2002a), but the axolotl spinal cord also regenerates after a more targeted transection injury (Butler and Ward, 1965, Butler and Ward, 1967, Clarke et al., 1988, O’Hara et al., 1992, Zukor et al., 2011). These lines of investigation have identified a population of Sox2^+^/GFAP^+^ glial cells that function as *bona fide* neural stem cells (NSCs) in the axolotl spinal cord (Echeverri and Tanaka, 2002, Fei et al., 2016, Fei et al., 2014a, McHedlishvili et al., 2012, Rodrigo Albors et al., 2015). These Sox2^+^/GFAP^+^ NSCs proliferate after injury and differentiate into new glia and neurons (Echeverri and Tanaka, 2002, McHedlishvili et al., 2012, Rodrigo Albors et al., 2015). Inhibition of NSC function by CRISPR/Cas9-mediated knockout of Sox2 results in deficient regenerative outgrowth of the spinal cord after tail amputation (Fei et al., 2016, Fei et al., 2014a).

Molecular signals required for the NSC response to injury have been identified after both tail amputation and spinal cord transection. Sonic hedgehog, Wnt/PCP, and Fgf signaling are indispensable for the pro-regenerative NSC response to tail amputation (Rodrigo Albors et al., 2015, Schnapp et al., 2005, Zhang et al., 2000, Zhang et al., 2002a). During spinal cord regeneration after transection, the transcriptional complex AP-1^cFos/JunB^ and MAP kinase signaling are critical regulators of the NSC response to injury (Sabin et al., 2015a, Sabin et al., 2019). Additionally, microRNA (miRNA) signaling is important to fine-tune the NSC response to injury after both tail amputation and spinal cord transection (Diaz Quiroz et al., 2014, Gearhart et al., 2015, Lepp and Carlone, 2014, Sehm et al., 2009).

We previously demonstrated that miR-200a is an important regulator of the glial cell response after spinal cord transection (Sabin et al., 2019). The function of miR-200a has been most extensively studied during neurodevelopment and epithelial-to-mesenchymal transition (EMT) (Trumbach and Prakash, 2015, Zaravinos, 2015). miR-200a functions to inhibit EMT by directly repressing the expression of the transcription factor *β-catenin* (Su et al., 2012, Zaravinos, 2015). This leads to maintained epithelial polarity and decreased Wnt signaling. During neurodevelopment, miR-200 family members regulate many processes: neuronal survival (Karres et al., 2007), neuroepithelial progenitor proliferation, NSC identity and neuroblast transition (Morante et al., 2013), neural progenitor identity and cell cycle dynamics (Peng et al., 2012), as well as fine tunes signaling networks necessary for neurogenesis (Choi et al., 2008, Vallejo et al., 2011) and gliogenesis (Buller et al., 2012).

Early experiments aimed at determining the potential of GFAP^+^/Sox2^+^ NSCs prospectively labeled these cells with the Glial fibrillary acidic protein (GFAP) promoter driving GFP expression and used live *in vivo* fluorescence imaging to follow GFP^+^ glial cells after tail amputation. Most GFP^+^ NSCs gave rise to new neurons and glia but a small proportion of labeled cells left the spinal cord and contributed to muscle and cartilage within the regenerated tail (Echeverri and Tanaka, 2002). Similar experiments using grafting of GFP^+^ spinal cords into non-transgenic animals further confirmed that spinal cord cells exited the spinal cord and contributed to cells of other lineages during tail regeneration (McHedlishvili et al., 2007).

Recent reports have identified a population of progenitors, called neuromesodermal progenitors (NMPs), which reside in the posterior of developing vertebrate embryos (Henrique et al., 2015, Kimelman, 2016b). NMPs are competent to contribute to both the mesoderm and spinal cord during embryonic development (Garriock et al., 2015, Henrique et al., 2015, Tzouanacou et al., 2009). Extensive genetic and biochemical analysis determined that NMPs can be defined by the co-expression of low levels of the transcription factors Brachyury and Sox2 (Gouti et al., 2017, Koch et al., 2017a, Turner et al., 2014a, Wymeersch et al., 2016). Also, the relative level of Fgf and Wnt signaling activity regulate NMP cell fate decisions (i.e. differentiation into mesodermal progenitors or neural progenitors) (Bouldin et al., 2015, Garriock et al., 2015, Goto et al., 2017, Gouti et al., 2017, Gouti et al., 2015, Martin, 2016, Martin and Kimelman, 2008, Turner et al., 2014a). Both Fgf and Wnt signaling are important regulators of the NSC response to tail amputation, as inhibition of either Wnt or Fgf blocks tail regeneration (Makanae et al., 2016, Ponomareva et al., 2015, Zhang et al., 2000, Albors et al., 2015). Although the role of individual Fgf ligands in spinal cord regeneration is relatively unknown, the expression of *wnt5* has been elegantly shown to be essential for orientated cell division and outgrowth of the spinal cord during whole tail regeneration (Albors et al., 2015). However, the activity of these pathways after spinal cord transection has not been well characterized.

In this study we identify a role for miR-200a in stabilizing the NSC identity after spinal cord transection in axolotl, by repressing expression of the mesodermal marker *brachyury*. Furthermore, we uncovered other genes in the miR-200 pathway and provide evidence that depending on the injury context, such as spinal cord lesion repair or spinal cord outgrowth during tail regeneration, that miR-200a plays an important role in determining the identity of NSCs in the spinal cord during the regenerative process.

## Results

### Transcriptional profiling identifies conserved miR-200a targets in homeostatic versus regenerating spinal cords

We previously identified miR-200a as a key microRNA (miRNA) that inhibits *c-jun* expression in neural stem cells in the spinal cord after injury, and hence plays an important role in preventing reactive gliosis and promoting a pro-regenerative response(Sabin et al., 2019). Additional RNA sequencing (RNA-seq) on uninjured and 4 days post injury spinal cord tissue electroporated with a control or specific, antisense miR-200a inhibitor identified other targets of miR-200a in axolotl spinal cord (**Fig. 1A**, **Table S1**). During normal regeneration at 4 days post injury there were 1,163 genes that are differentially expressed (Log2 fold change ≥2-fold, p≤0.05) compared to uninjured spinal cords. Inhibition of miR-200a in the uninjured spinal cord resulted in 6,235 transcripts with a greater than 2-fold differential expression compared to control uninjured spinal cords. Interestingly, of the 6,235 differentially expressed genes, only 2,760 were up-regulated (**Fig.1A**, **Fig. S1**, **Table S1**). We used GOrilla analysis to identify gene ontology (GO) terms for the subset of genes that were significantly up-regulated after miR-200a inhibition. GO terms involved with translation, RNA metabolism, peptide metabolism, and translation initiation were significantly enriched in this gene set (p≤10^-24^) (**Fig.S1**). Interestingly, the 3,475 genes that were significantly down-regulated in uninjured spinal cords after miR-200a inhibition were enriched for GO terms involved with organismal development, developmental process, cellular differentiation, and signaling (p≤10^-22^) (**Fig. S1**).

**Figure 1.**
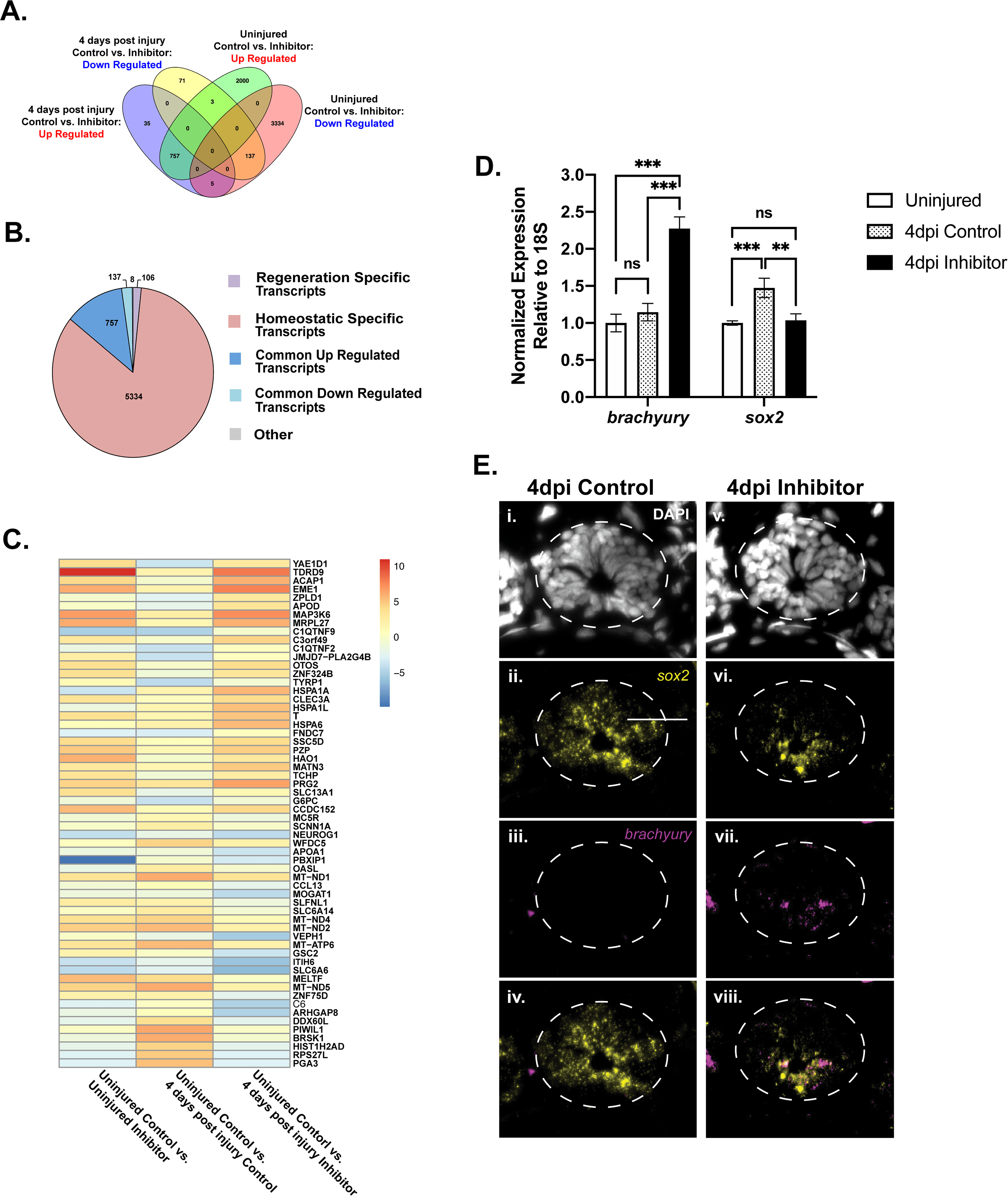
miR-200a inhibition during spinal cord injury leads to *brachyury* expression axolotl in spinal cord stem cells. (A) RNA-Sequencing analysis identified a large subset of differentially regulated genes following injury. The Vehn diagram compares the number of overlapping differentially expressed genes between uninjured, 4 days post injury control and miR-200a inhibitor treated samples. (B) Pie-chart showing the relative portion of all transcripts that are differentially regulated. Regeneration specific transcripts are defined as differentially expressed transcripts with a log2fc > 1 or < 1 and padj <0.05 between 4dpi control animals and 4dpi animals treated with mir-200a inhibitor that were not differentially expressed in uninjured animals. (C) Log2fold change heat map demonstrates the 30 most up- and down-regulated genes in uninjured and 4 days post injury control versus miR-200a inhibitor treated spinal cords. This analysis revealed that the transcription factor *brachyury* (*T*) is dramatically up-regulated after miR-200a inhibition. (D) qRT-PCR analysis confirmed that miR-200a inhibition led to increased *brachyury* expression and blocked the up-regulation of the neural stem cell marker *sox2* in 4 days post injury (dpi) spinal cords (n=3). (E) Fluorescent *In situ* hybridization confirmed the qRT-PCR analysis and revealed miR-200a inhibition leads to *brachyury* expression (n=5) in spinal cord cells and a failure to up-regulate *sox2* expression (n=3). *p≤0.05, **p≤0.01, *** p≤0.001, N.S. is not significant. Error bars represent ±S.T.D. Scale bar= 50μm.

Analysis of differentially expressed genes at 4 days post injury after miR-200a inhibition identified a total of 1,007 genes that were differentially expressed compared to control spinal cords. This is a much smaller gene set and suggests that there is a higher specificity to genes affected by miR-200a during spinal cord regeneration. A total of 797 genes were up-regulated and 210 genes were down-regulated after miR-200a inhibition (**Fig. S1**). Genes that were up-regulated were enriched for GO terms involved with nucleic acid metabolism, specifically RNA metabolism, and protein localization (p≤10^-6^) (**Fig. S1**). Interestingly, the top GO terms enriched in down-regulated genes were involved with nervous system processes, specifically synaptic signaling and chemical synaptic signaling, as well as nervous system development (p≤10^-6^).

Taking a more targeted gene-level approach, we generated a heat map of the 30 most significantly up-regulated and down-regulated genes in all four conditions (**Fig.1C**). Consistent with the GO analysis, genes involved in RNA processing, nucleic acid metabolism, and protein targeting were among the most up-regulated genes in our data set (*tdrd9*, *acap1*, *eme1*, *zfp324b*). Similarly, genes involved with neurotransmitter transport, neuronal polarization, neurotrophin signaling, and neuronal differentiation were among the most down-regulated genes (*slc6a6*, *brsk1*, *slc6a14*, *arhgap8*, *neurog1*). Surprisingly, the transcription factor *brachyury* (*T*) was among one of the most up-regulated genes at 4 days post injury in response to miR-200a inhibition (**Fig.1C**). In 4 days post injury controls, *brachyury* was not up-regulated in response to injury, the RNA seq transcripts per million (TPM) values on Control uninjured were 0.782 TPM, versus 4 dpi post injury 0.92 TPM, show no significant increase (**Table S1**). However, inhibition of miR-200a in uninjured spinal cords led to a 2-fold increase in *brachyur* expression (2.236 TPM), while the combination of miR-200a inhibition during injury led to a highly significant 7-fold increase in its mRNA levels (5.8 TPM, **Table S1**).

We used quantitative RT-PCR (qRT-PCR) to verify genes of interest revealed by RNA-seq, this approach confirmed that *brachyury* is detectable at very low levels in uninjured and control 4 days post injury spinal cords but is significantly up-regulated after miR-200a inhibition in 4 days post injury spinal cords (**Fig.1D**). This is an intriguing finding as *brachyury* is considered a classical marker of mesodermal tissue and was originally thought to be absent from in the nervous system. However, more recent research has identified a bipotent cell population in development, in which some spinal cord neural progenitor cells are developmentally derived from Sox2^+^/Brachyury^+^ neuromesodermal progenitors (NMPs) (Garriock et al., 2015, Tzouanacou et al., 2009, Wymeersch et al., 2016). In the axolotl, the *bona fide* stem cells that line the central canal are identified by the expression of the glial cell marker GFAP and the neural stem cell marker Sox2. These GFAP^+^/Sox2^+^ cells respond to the injury, divide, migrate and repair a lesion in the spinal cord, or regenerate lost cells and tissues in the context of whole tail regeneration (Sabin et al., 2015a, Fei et al., 2014b, Echeverri and Tanaka, 2002, Echeverri and Tanaka, 2003a, McHedlishvili et al., 2007, McHedlishvili et al., 2012). Given that NMPs and axolotl glial cells both express Sox2 and that *sox2* is a miR-200a target during mouse brain development (Peng et al., 2012), we assayed *sox2* transcript abundance. Interestingly, while *sox2* is slightly up-regulated in control 4 days post injury compared to uninjured spinal cords, *sox2* expression did not increase in miR-200a inhibitor treated spinal cords. Instead, the *sox2* transcript abundance remains near uninjured homeostatic levels (**Fig.1D**). This observation suggests that axolotl *sox2* is not a direct target of miR-200a as it is in mammals (Pandey et al., 2015, Peng et al., 2012, Wang et al., 2013).

To identify the cells that express *brachyury* in the 4 days post injury spinal cord after miR-200a inhibition, *in situ* hybridization was used. *In situ* hybridization determined that cells lining the central canal are *brachyury^+^* after miR-200a inhibition (**Fig.1Evii**), and importantly, this is the same population of cells that express *sox2 (***Fig.1Evi, viii**). Collectively, this data indicates that miR-200a inhibition leads to increased *brachyury* expression in stem cells in the axolotl spinal cord. Although these progenitor cells have been traditionally thought of as NSCs due to their expression of the classical NSC marker *sox2,* they also express low levels of the mesodermal marker *brachyury* (**Fig.1D**), suggesting that they are in fact a population of bipotent stem cells and may in fact have broader differentiation potential.

### Inhibition of miR-200a leads to changes in cell fate after spinal cord injury

To determine if miR-200a inhibition and subsequent upregulation of the mesodermal marker *brachyury* leads to changes in the number of NSC in the regenerating spinal cord, we quantified the number of Sox2^+^ cells in control versus inhibitor treated animals **(Fig.2A, B**). Previous work has shown that after spinal cord injury the cells that are activated to partake in the regeneration process lie within 500μm of the injury site. However, we detected no significant difference in the total number of Sox2^+^ cells 500μm rostral or caudal of the injury site **(Fig.2A, B)**. During the normal regenerative process in the spinal cord, the Sox2^+^ stem cells will replenish the GFAP^+^ cell population and differentiate into new neurons (Albors et al., 2015, Echeverri and Tanaka, 2002, Echeverri and Tanaka, 2003a, McHedlishvili et al., 2007, McHedlishvili et al., 2012). To determine if the number of newborn neurons is affected by miR-200a inhibition we quantified the number of NeuN and EDU^+^ cells 500μm rostral and caudal to the injury site. We found that overall, significantly fewer NeuN^+^/EdU^+^ cells were found in the miR-200a inhibitor animals **(Fig. 2C, D).** Collectively, these data demonstrated that the total number of the Sox2^+^ spinal cord stem cells is relatively the same in control versus miR-200a inhibited animals, but less NeuN^+^ cells are found in the inhibitor treated animals. These findings suggest that either more cells remain in a progenitor-like state, or the expression of *brachyury* in the miR-200a inhibitor treated animals changes the fate of the cells. To address this question, we used *in vivo* cell tracking to determine the fate of these cells during regeneration of the lesioned spinal cord. We have previously tracked the fate of GFAP^+^ spinal cord stem cells during regeneration of the lesioned spinal cord and found that these cells proliferate and migrate to replace the portion of injured neural tube, and that this is a bidirectional process (Sabin et al., 2015a). The same technique was used in these studies, the axolotl GFAP promoter driving expression of a fluorescent protein was injected into the lumen of the spinal cord, the animals were electroporated to label small groups of cells. The miR-200a inhibitor was injected into animals with fluorescently labelled cells and then the spinal cord ablation was performed (Sabin et al., 2019). The animals were imaged every 3-days over a two-week time period. In the control labelled animals, we found the cells behaved as we had previously described, the labeled cells proliferated and partook in repair of the neural tube, replenished the endogenous stem cell population, and differentiated to replace lost neurons (**Fig.3** **A-D),** consistent with our previous work (Sabin et al., 2015). In contrast, in the miR-200a inhibitor treated animals although the cells proliferated and partook in repair of the lesioned spinal cord, we also discovered that the cells exited from the spinal cord and differentiated into muscle cells. The labelled cells which started in the spinal cord where always found in the muscle layer adjacent or directly above the spinal cord (**Fig.3I, J**). In all miR-200a inhibitor treated animals we observed at least 1 muscle fiber being formed in all animals (n=25), although in some animal’s multiple fibers were seen. Additionally, in inhibitor and control animals’ some cells differentiated into neurons and remained within the neural tube to give rise to new glial cells (data not shown). This data suggests that miR-200a represses *brachyury* in *sox2*^+^ spinal cord stem cells, maintaining the cells in neural primed state. Inhibition of miR-200a in these cells results in the co-expression of neural (*sox2*) and mesoderm (*brachyury*) markers, converting the cells into a bipotent progenitor population capable of making both neural and mesodermal cells.

**Figure 2:**
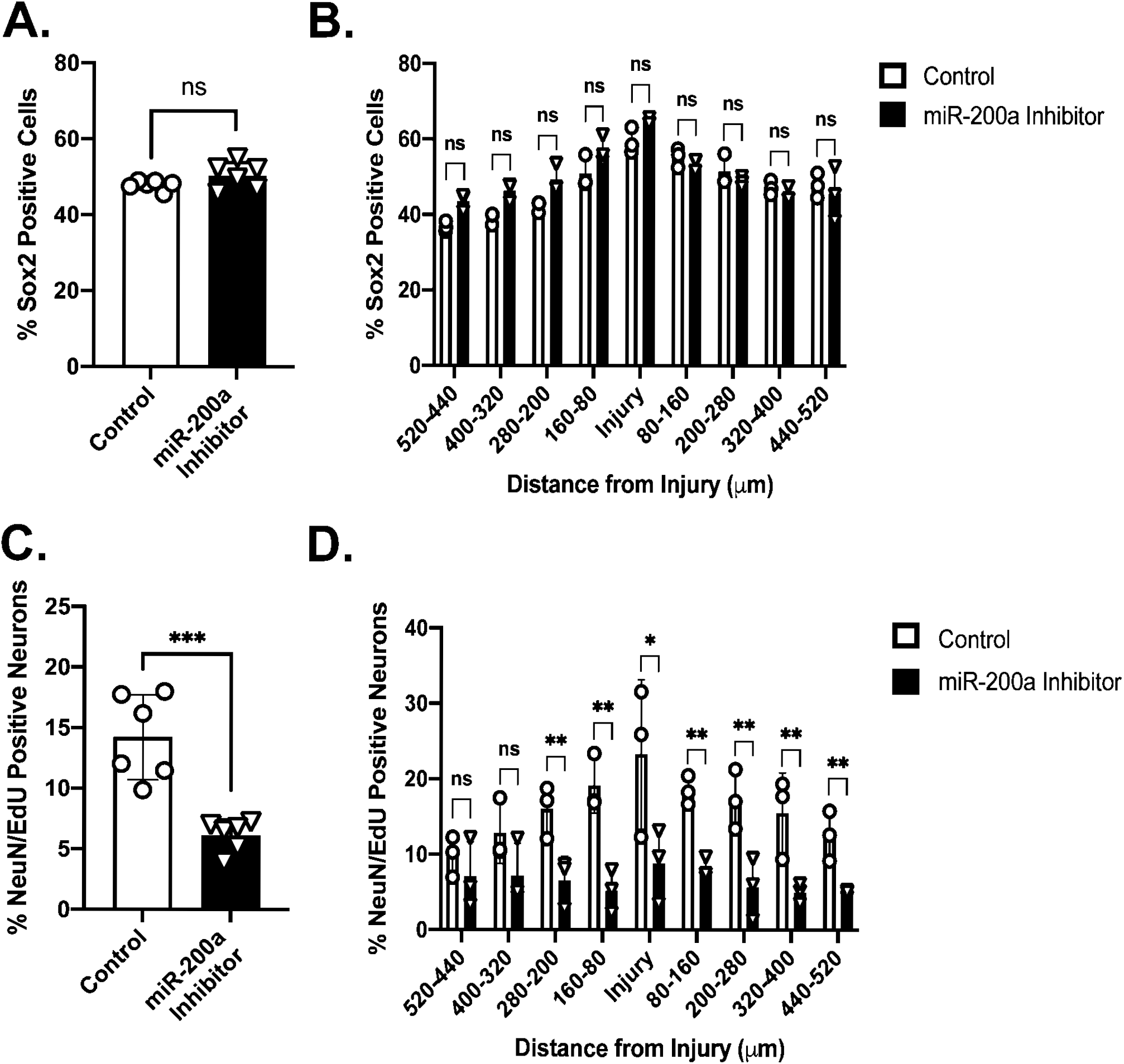
Chronic miR-200a inhibition affects the birth of new neurons. Inhibition of miR-200a for 2 weeks does not affect the (A) proportion or (B) distribution of Sox2^+^ stem cells in the spinal cord throughout the regeneration zone (n=6). Inhibition of miR-200a for 2 weeks results in an smaller proportion of new born neurons compared to controls (C). (D) An increased proportion of new born neurons reside closer to the injury site while chronic miR-200a inhibition blocked this increase in new born neurons close to the injury site. (n=6). *p≤0.05, **p≤0.01, *** p≤0.001, N.S. is not significant. Error bars represent ±S.T.D.

**Figure 3:**
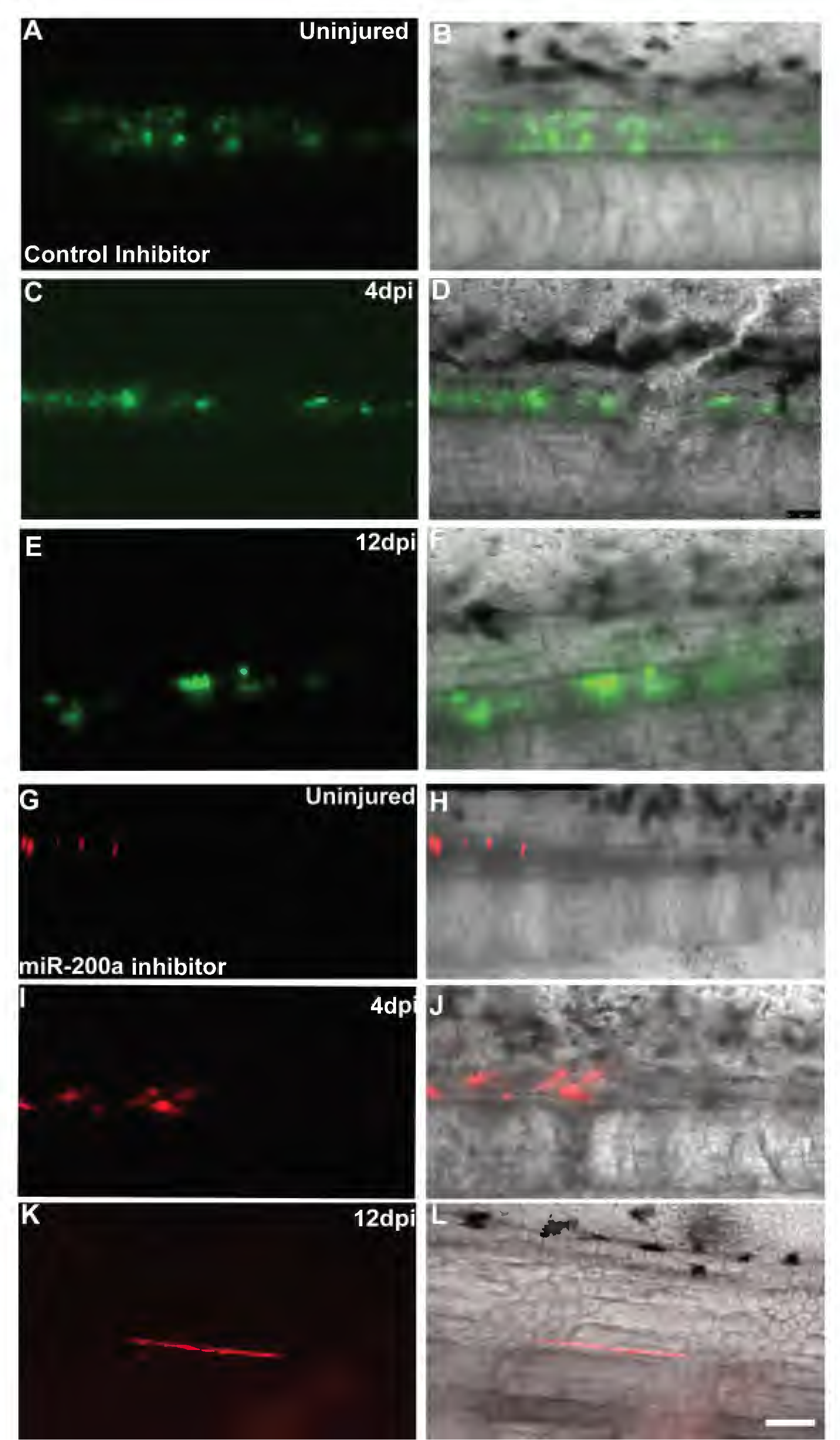
miR-200a inhibited spinal cord cells form muscle during spinal cord lesion repair. The cells lining the central canal of the spinal cord were labelled using GFAP promoter driving GFP or tdTomato by injection and electroporation. Control cells were followed over a 14 day period and cells gave rise to new glial cells or neurons only (A-F) (n=20). Cells which were injected with the miR-200a inhibitor were followed in parallel over the same time period and were found to exit the spinal cord and give rise to muscle cells (G-L). (n=25). Scale bar = 50μm.

### Molecular regulation of progenitor cells by miR-200a

Our data strongly indicate that inhibition of miR-200a leads to the expression of *brachyury* in stem cells within the axolotl spinal cord (**Fig. 1D**,). However, the signaling pathway(s) upstream of *brachyury* expression were not known. As a first step we first tested whether miR-200a could directly repress *brachyury* expression. The axolotl *brachyury* 3’ untranslated region (UTR) contains three miR-200a seed sequences, this indicates that miR-200a could directly regulate *brachyury* expression. Consistent with this hypothesis, co-transfection of B35 neural cells with a *brachyury* 3’ UTR luciferase reporter and miR-200a mimic led to decreased luciferase activity compared to the control mimic (**Fig. S2A**). This finding confirmed that miR-200a directly represses *brachyury* expression in axolotl spinal cord stem cells in homeostatic conditions and during normal regeneration.

During normal spinal cord regeneration in the context of a tail amputation model it has been found that *wnt* genes are re-expressed in the caudal 500μm of the outgrowing spinal cord and are necessary for this outgrowth (Albors et al., 2015).Further studies have shown that inhibition of all Wnt or Fgf signaling after tail amputation abolished regenerative outgrowth, suggesting both are necessary for spinal cord and tail regeneration (Ponomareva et al., 2015). As both Fgf and Wnt signaling regulate cell fate decisions of brachyury^+^/sox2^+^ NMPs during development, we first tested whether Fgf signaling could be affected by miR-200a inhibition during regeneration. We assayed for expression of *fgf8* and *fgf10* by qRT-PCR on isolated spinal cord tissue. *Fgf8* expression was slightly down-regulated at 4 days post injury after miR-200a inhibition compared to uninjured spinal cords (**Fig.4A**), while *fgf10* expression was significantly up-regulated after miR-200a inhibition in 4 days post injury spinal cords compared to controls (**Fig.4A**). This finding is consistent with the idea that miR-200a inhibition could lead to an increase in *fgf* ligand expression in regenerating spinal cords. However, given that Wnt signaling directly regulates Brachyury expression (Arnold et al., 2000, Yamaguchi et al., 1999) and NMP cell fate decisions (Bouldin et al., 2015, Garriock et al., 2015, Martin, 2016, Martin and Kimelman, 2008), we wanted to further examine the role of Wnt signaling.

**Figure 4:**
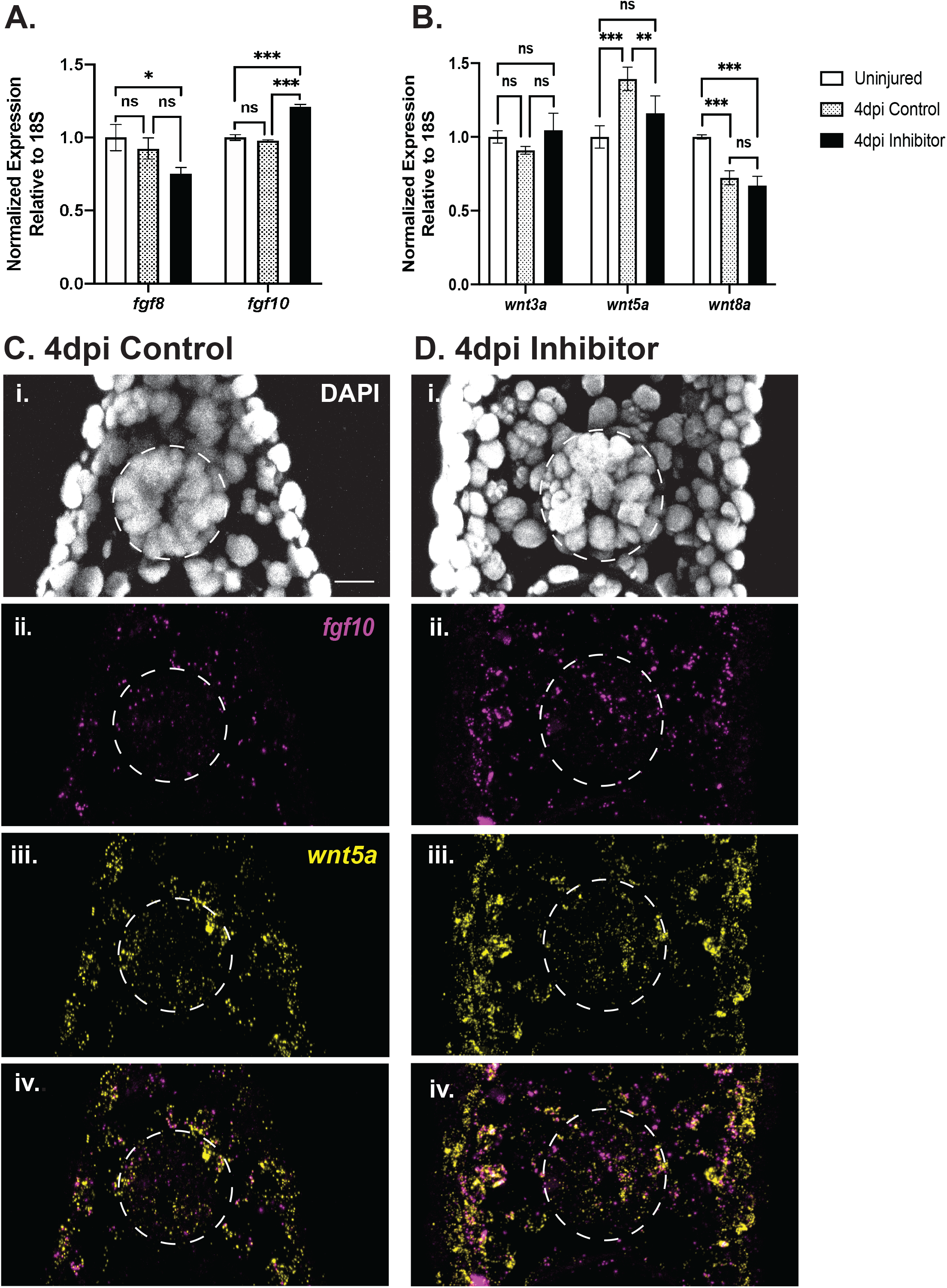
miR-200a inhibition affects the expression of Wnt and FGF signaling ligands. (A) qRT-PCR analysis revealed fgf10, but not *fgf8,* was significantly up-regulated after miR-200a inhibition (n=3). (B) qRT-PCR analysis showed that miR-200a inhibition differentially affects the expression of *wnt5a*, but not additional Wnt ligands (*wnt3a*, *wnt8a*; n=3). (C) Fluorescent *in situ* hybridization in control (C) and miR-200a inhibitor treated (D) animals confirmed the qRT-PCR analysis, and demonstrates an increase in *fgf10* expression and a down-regulation of *wnt5a* within stem cells in the spinal cord (n=6). *p≤0.05, **p≤0.01, *** p≤0.001, N.S. is not significant. Error bars represent ±S.T.D. Scale bar= 50μm.

The expression levels of *wnt3a, wnt5a, wnt8a* were quantified using qRT-PCR (**Fig. 4B**). Both *wnt3a* and *wnt8* transcript levels were not significantly altered in the inhibitor treated animals compared to controls. However, we did detect a significant difference in *wnt5a* levels. In 4 day post injury controls, *wnt5a* was up-regulated after injury, although this change in expression was not found in the miR-200a inhibitor treated animals. To further verify the qRT-PCR results for *fgf* and *wnt* genes we performed fluorescent *in situs* for *fgf10* and *wnt5*a in control and miR-200a inhibitor treated regenerating animals. This confirmed that indeed *fgf10* transcript levels are up-regulated in cells within the spinal cord in comparison to the control regenerating animals (**Fig.4C, D**). Additionally, *wnt5a* transcript levels were down-regulated in cells within the spinal cord in comparison to controls (**Fig.4C, D**). Although here we see only changes in *wnt5a* expression, a Wnt which is known to play an important role in regeneration (Albors et al., 2015), there are many additional Wnt ligands, therefore Wnt signaling activity could still be affected by miR-200a inhibition. To establish a baseline for Wnt signaling activity after spinal cord injury we assayed *lef1* expression, which is a direct transcriptional target downstream of Wnt signaling (Filali et al., 2002). *Lef1* expression was significantly up-regulated in control 4 days post injury compared to uninjured spinal cords (**Fig. S3A**), indicating a potential increase in Wnt signaling after injury. Remarkably, *lef1* expression was significantly up-regulated even further after miR-200a inhibition in 4 days post injury compared to control regenerating spinal cords **(Fig. S3A**). Collectively, these data indicate that miR-200a inhibition could result in increased Wnt signaling, potentially independent of changes in *wnt* ligand expression.

### miR-200a modulates Wnt signaling activity by directly targeting β-catenin

While miR-200a inhibition could lead to increased Wnt signaling, it was not clear how this was occurring. During tumor progression, miR-200a inhibits the epithelial-to-mesenchymal transition subsequently blocking tumor cell metastasis (Su et al., 2012, Zaravinos, 2015). This is partially achieved through the direct repression of *β-catenin* by miR-200a, resulting in decreased Wnt signaling (Su et al., 2012). We did not observe a significant up-regulation of specific *wnt* ligand expression after miR-200a inhibition in the spinal cord cells (**Fig. 4B**). However, as determined by *lef1* expression, miR-200a inhibition could lead to increased Wnt signaling (**Fig. S3A**). Therefore, we hypothesized that miR-200a might regulate Wnt signaling by targeting *β-catenin*. To test our hypothesis, we first assayed for changes in *β-catenin* transcript abundance (*ctnnb1*). qRT-PCR analysis confirmed that after injury in control 4 days post injury spinal cords, there is an increase in *ctnnb1* abundance compared to uninjured spinal cords, similar to what we observed for *lef1* (**Fig. S3A**). There is a slight increase of *ctnnb1* transcript levels after miR-200a inhibition compared to control 4 days post injury spinal cords (**Fig. S3A**), indicating *β-catenin* could be a direct target of miR-200a in axolotl.

To determine whether miR-200a could target axolotl *β-catenin*, we cloned the *ctnnb1* 3’ UTR and identified two miR-200a seed sequences. We subcloned the *ctnnb1* 3’ UTR into a luciferase reporter and co-transfected cells with a control mimic or miR-200a specific mimic. There was decreased luciferase activity in miR-200a mimic transfected cells compared to control, suggesting that miR-200a could regulate *β-catenin* expression (**Fig. S3B**). To confirm that the decrease in luciferase activity is due to direct regulation by miR-200a, we mutated both seed sequences in the *ctnnb1* 3’ UTR and repeated the luciferase experiments. Mutation of the miR-200a seed sequences completely alleviated the repression, confirming that axolotl *β-catenin* is a direct target of miR-200a, similar to mammals (**Fig. S3B**).

Taken together, these data are consistent with the idea that miR-200a could modulate Wnt signaling through the direct regulation of *β-catenin* transcript levels. Inhibition of miR-200a leads to increased *lef1* expression, which is indicative of increased Wnt signaling. Increased levels of Wnt signaling may contribute to the increased *brachyury* expression and changes in *fgf10* levels in axolotl stem cells after spinal cord lesion.

### The role of spinal cord stem cells in spinal cord injury versus tail regeneration

We have shown that when a lesion occurs in the axolotl spinal cord the glial cells adjacent to the injury site respond to the injury cue and proceed to behave like NSCs; they divide, migrate, self-renew and replace lost neurons. However, previous work has shown that during spinal cord regeneration after amputation, rather than injury, these glial cells can transdifferentiate and give rise to cells of both the ectodermal and mesodermal lineage (Echeverri and Tanaka, 2002, McHedlishvili et al., 2007). We next examined the expression of *brachyury* in the context of whole tail regeneration and discovered that *brachyury* is expressed in the *sox2^+^* stem cells of the spinal cord 500μm adjacent to the injury site at 7-days post amputation (**Fig. 5**). To determine if this is an attribute of the larval animals only, we also examined regenerating tail tissue from 2-year old adult animals and found that in response to tail amputation that these progenitor cells in the adult spinal cord indeed co-express *brachyury* and *sox2* during tail regeneration in adult (**Fig. 5**) and larval animals (**Fig.S4**). These data suggest that during the regenerative process the cells lining the central canal determine what tissue types need to be restored. When only a small portion of the neural tube must be regenerated following injury, the progenitor cells adopt a neural stem cell state to successfully regenerate the spinal cord. During whole tail regeneration following amputation when multiple tissue lineages must be regenerated, these cells within the spinal cord become bipotent progenitors capable of making mesoderm and ectoderm (**Fig.6**). Collectively, these experiments have shed light on the context dependent nature of miRNA signaling during spinal cord lesion repair versus tail amputation and have identified new signaling pathways that regulate progenitor cell fate during axolotl spinal cord regeneration.

**Figure 5.**
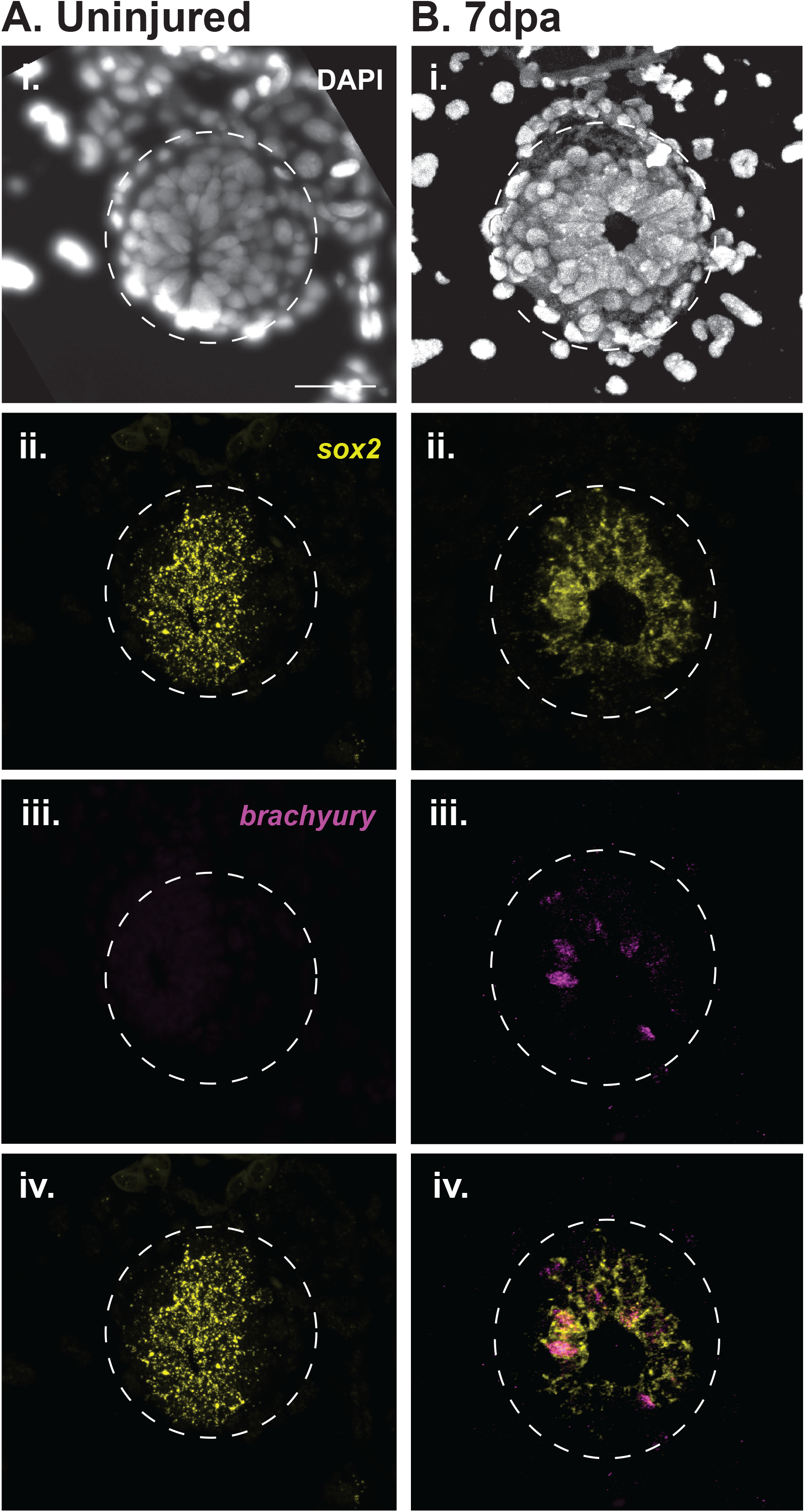
Spinal cord amputation leads to *brachyury* expression in spinal cord stem cells. (Ai-iV) Fluorescent *in situ* hybridization revealed that *sox2* is abundant within the uninjured adult spinal cord, while *brachyury* is absent (n=2). (Bi-iv) At 7 days post amputation, *brachyury* was localized to spinal cord stem cells that share an overlapping expression pattern with *sox2* (n=2). Scale bar= 50μm.

**Figure 6.**
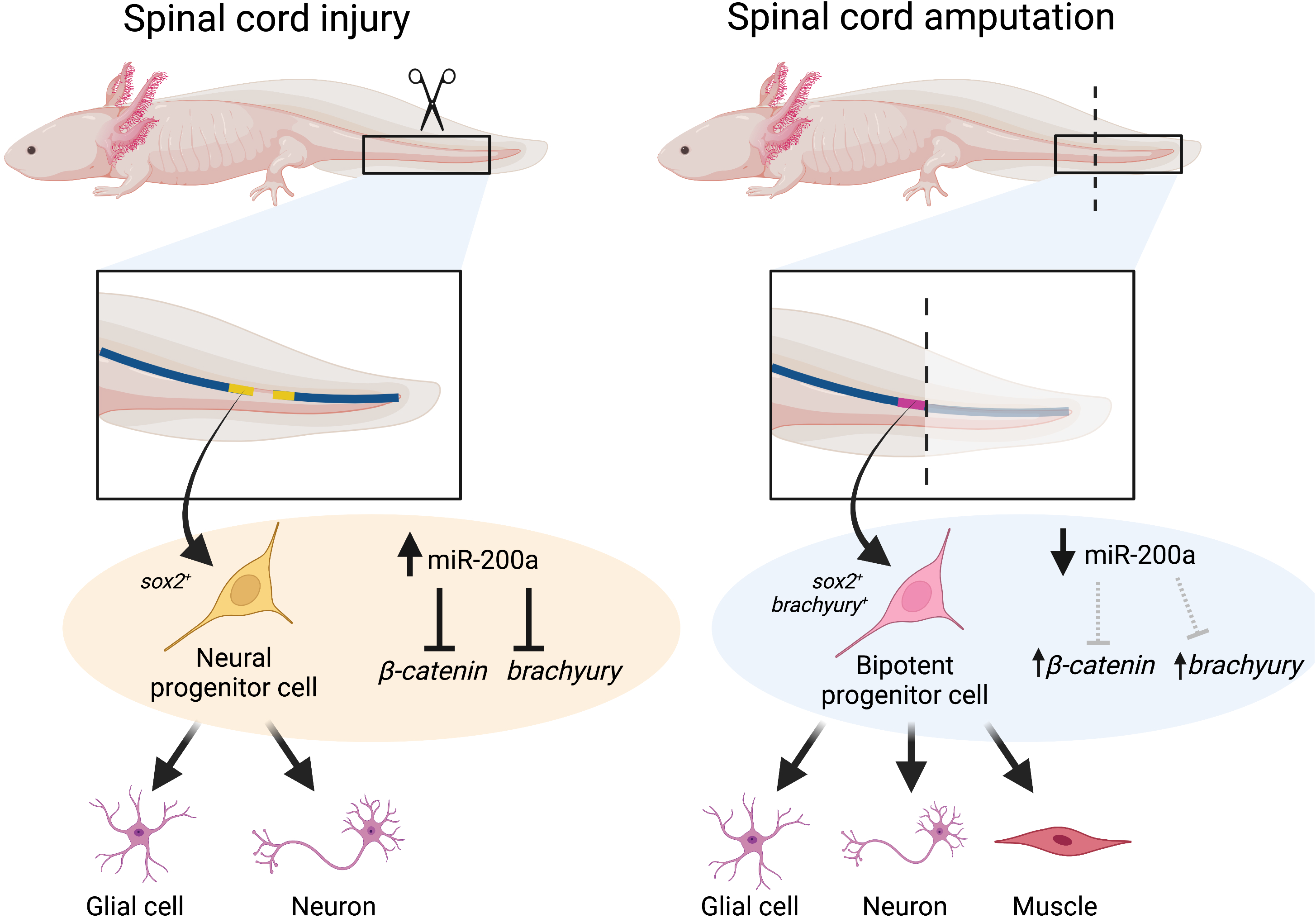
A proposed model for the role that miR-200a plays in different injury paradigms. When a lesion occurs in the spinal cord miR-200a levels remain high, which inhibits *brachyury* expression and modifies levels of *β-catenin*, potentially stabilizing a neural stem cell identity in the cells adjacent to the injury site. After spinal cord injury these cells replace neurons and glial only. In contrast, when the tail is amputated progenitor cells respond to injury cues and replace multiple cell types of different developmental origins. These cells in the spinal cord then up-regulate *brachyury* in the *sox2*^+^ stem cells of the spinal cord and direct these cells to proliferate and form cells of both ectodermal and mesodermal origin.

## Discussion

The current study has identified miR-200a as a regulator of stem cell fate in the regenerating axolotl spinal cord. GO term analysis of genes down-regulated in the uninjured and 4 days post injury spinal cord after miR-200a inhibition showed that these genes were involved with nervous system development, organismal development, synaptic signaling, and cellular differentiation (**Fig.1, Fig.S1**). Specifically, genes involved with neuronal differentiation (*neurog1, neuroD4*) and neuronal processes like synaptic transmission (*CHRNB1, GABRA4*) and neurotransmitter uptake (*SLC6A6, SLC18A3, SLC6A14*) were down-regulated (**Fig.1**, **Fig. S1**). This suggests that miR-200a normally functions to promote NSC identity. This is consistent with multiple reports across various species that inhibition of miR-200a and other miR-200 family members results in the loss of neural progenitor identity and precocious neuronal or glial differentiation (Buller et al., 2012, Choi et al., 2008, Morante et al., 2013, Peng et al., 2012, Trumbach and Prakash, 2015, Vallejo et al., 2011). However, we have found that in the axolotl spinal cord that even in uninjured conditions in larval or adult axolotls, the cells lining the central canal in fact express low levels *brachyury* and *sox2*, the classical markers of mesoderm and neural stem cells (**Fig.1D**, **Fig.5**). These cells may represent a bipotent progenitor cell population and our data suggests that increased levels of *brachyury* are necessary for a progenitor to make the decision to exit the spinal cord and become a cell type of mesodermal origin (**Fig.6**).

During embryonic development in multiple species a small population of cells that co-express Sox2 and Brachyury have been identified and are now called neuromesodermal progenitor cells (Gouti et al., 2014, Henrique et al., 2015, Jurberg et al., 2013, Kimelman, 2016b, Taniguchi et al., 2017, Tsakiridis et al., 2014, Tsakiridis and Wilson, 2015, Turner et al., 2014b, Tzouanacou et al., 2009). Neuromesodermal progenitor cell commitment to the neural lineage is partially determined by the relative levels of Sox2 compared to Brachyury, given that the two transcription factors function to antagonize one another (Koch et al., 2017a) and by the respective levels of *fgf* versus *wnt* that the progenitor cells encounter (refs). During axolotl development to date a definitive population of neuromesodermal progenitors has not been defined, however work published by Taniguichi et al has shown that a posterior region of the axolotl neural plate is positive for *brachyury* and *sox2* and this region gives rise to mesoderm during development (Taniguchi et al., 2017).This finding is consistent with the idea that the axolotl may also have a bipotent progenitor pool of cells established during early development, however more work is needed, especially lineage tracing to establish if these cells behave similar to what is seen in other species like chick, mouse and zebrafish. Our results here showing that by qRT-PCR and RNAscope *in situs* that *brachyury* and *sox2* are detected in the progenitor cells of the spinal cord in both larval and adult axolotls would suggest that axolotls retain a population of cells in the spinal cord throughout life that are bipotent. Work from McHedlishvili et al previously showed that adult axolotl retains expression of embryonic markers of dorsal/ventral patterning like *pax7*, *pax6* and *shh* genes that are not expressed in adult mammalian spinal cord (McHedlishvili et al., 2007). They additionally showed that like earlier lineage tracing work in axolotl, cells from the spinal cord do in fact migrate out and form a range of other cell types including blood vessels, skin, cartilage, and muscle cells. Overall, these bodies of work indicate that the cells in the axolotl spinal cord retain a multi-potent progenitor cell state and are capable to respond to injury cues which direct them towards different cell fates as needed. Very early work on tail regeneration in salamanders had already hinted that the terminal vesicle structure formed at the growing end of the spinal cord during tail regeneration is an area of epithelial to mesenchymal transition where cells delaminate from the neural tube and exit to contribute to regeneration of surrounding tissues of other developmental lineages (Benraiss et al., 1997, Egar and Singer, 1972, O’Hara et al., 1992). Our data now gives molecular insights into the identity of these cells. We found that miR-200a inhibited cells increase levels of *brachyury* and then these cells during regeneration of a spinal cord lesion will form muscle which is not observed in control regenerating lesions. However, we have not observed these cells to form cartilage, skin, fin mesenchyme of any other cell type. We cannot rule out that they have this potential, but the lineage tracing is limited as the fluorescent protein expression is driven by the GFAP promoter, we expect this promoter is turned off as the cells differentiate and as we can only image very 3 days, we may in fact miss some differentiation events. We observed in all animals where miR-200a is inhibited that at least one muscle fiber is formed from the labelled cells, however, again we may be missing some differentiation events due to the limitations of our labelling technique.

It is still not clear whether Brachyury directly regulates Sox2 levels in the regenerating axolotl spinal cord or whether it is via an indirect mechanism. Work from other labs in other research organisms has indicated that Brachyury and Sox2 can have a mutually repressive relationship (Kimelman, 2016a, Koch et al., 2017b, Martin, 2016). We have shown that miR-200a directly regulates *brachyury* and *β-catenin* via seed sequences in the 3’UTR of these genes. When miR-200a is inhibited in the spinal cord cells, *brachyury* is expressed at higher levels in these cells but *fgf* and *wnt* levels are also perturbed.

Work in development on NMP’s has shown that feedback loops exist between *brachyury, fgf and wnt* genes, and hence a complex signaling network may exist that is driven by specific levels of certain regulators in these cells at particular times. An additional level of complexity is the fact that Wnt is a secreted protein and although we see downregulation of it within the progenitor cells in the spinal cord, we also see that cells outside the spinal cord express *wnt* (**Fig.4**) and therefore the progenitor cells may also be influenced by external gradients of Wnt protein.

During development, Wnt and Fgf signaling tightly regulate neuromesodermal cell fate decisions (Goto et al., 2017, Gouti et al., 2017, Gouti et al., 2015, Martin, 2016) and both genes are known to play important roles in regeneration (Sun et al., 2002, Wilson et al., 2000, Zhang et al., 2000, Zhang et al., 2002b, Caubit et al., 1997, Ghosh et al., 2008, Lin G, 2008, Stoick-Cooper et al., 2007, Tanaka and Weidinger, 2008, Wehner et al., 2017, Zakany and Duboule, 1993). Canonical Wnt signaling is crucial for radial glial cell proliferation during neural tube development (Shtutman et al., 1999) and spinal cord regeneration in zebrafish (Briona et al., 2015). Therefore, it is not surprising to see a potential increase in Wnt signaling during spinal cord regeneration in axolotl. However, it is interesting that miR-200a does not regulate expression of *wnt* ligands, but instead regulates *β-catenin* levels (**Fig. S3**). This is reminiscent to the role of miR-200a in inhibiting EMT by repressing *β-catenin* and canonical Wnt signaling (Su et al., 2012, Zaravinos, 2015). The increase in *ctnnb1* levels after miR-200a inhibition is not statistically significant, however slight changes in transcript abundance can have profound effects on protein levels (Schwanhausser et al., 2011). Therefore, a modest increase in transcript abundance could represent a biologically significant increase in *β-catenin* protein levels.

The signals that inform injured cells what tissue must be replaced remain a mystery. Here we show that glial cells in the spinal cord appear to sense the difference between a lesion of the spinal cord that primarily needs replacement of neural stem cell and neurons, versus regeneration in the context of whole tail regeneration where cells of multiple developmental germ layer origin must be regenerated. Interestingly we find that cells of both the larval and adult tail regenerate bipotent progenitors that express *brachyury* and *sox2* in response to tail amputation, suggesting that the presence of these bipotent progenitors is not only a hallmark of embryonic development, but rather a stem cell population that is maintained in the animals specifically for regeneration. In the future it will be important to determine if all cells in the spinal cord have this potential or whether there are sub-populations of stem cells present in the axolotl spinal cord.

## Materials and Methods

### Animal Handling and Spinal Cord Injury

All axolotls used in these experiments were obtained and bred at the University of Minnesota or the Marine Biological Laboratory in accordance with IACUAC regulations. Prior to all *in vivo* experiments, animals (3-5cm) were anesthetized in 0.01% p-amino benzocaine (Sigma). Spinal cord ablations were performed as previously described (Diaz Quiroz et al., 2014,Sabin et al., 2015a). Briefly, a 26-gage needle was used to clear away skin and muscle to expose the spinal cord 6-10 muscle bundles caudle to the cloaca. Then, using the needle, a segment of spinal cord 1 muscle bundle thick, approximately 500μm, was removed. Animals were placed in cups and monitored for the duration of the experiments.

### Immunohistochemistry and EdU

Tissue was harvested and fixed in fresh 4% paraformaldehyde (Sigma) overnight at 4°C. Then tails were washed three times in phosphate buffered saline + 0.1% Tween 20 (PBSTw). Next the tails were incubated in a 50:50 solution of PBSTw and 30% sucrose. Finally, tails were transferred to 30% sucrose solution and allowed to equilibrate overnight at 4°C. The next day samples were embedded for cross-sectioning in TissueTek (Sakura) and stored at -20°C.

For EdU staining, animals were injected intraperitoneal with EdU at a concentration of 0.5 μg/μL in PBS+1% Fast Green at 5 and 7-days post injury then harvested at 14-days post injury. The tissue was processed for sectioning as described above and stained using the Click-iT EdU Imaging Kit (Invitrogen) according to the manufacturer’s instructions.

After staining for EdU, samples were processed for immunohistochemical analysis using either anti-Sox2 (Abcam) or anti-NeuN (Chemicon) primary antibodies as previously described (Sabin et al., 2015a, Sabin et al., 2019). Briefly, slides were subjected to a boiling citrate antigen retrieval step and then washed with PBSTw 3 times for 5 minutes each. Samples were blocked (PBS+0.1% Triton-X+2% bovine serum albumin +2% goat serum) for an hour at room then incubated overnight at 4°C in primary antibodies diluted (1:100) in blocking buffer. The next day, slides were washed 4 times with PBSTw and then incubated with secondary antibody (Invitrogen) diluted in blocking buffer (1:200) for 2 hours at room temp and cell nuclei were counterstained with 4’,6-diamidino-2-phenylindole (DAPI) (1:10,000). After secondary incubation the slides were washed four times with PBSTw and mounted in Prolong Anti-fade mounting solution (Invitrogen). All samples were imaged using an inverted Leica DMI 6000B fluorescent microscope.

All images were generated using Fiji and cells were counted with the Cell Counter plugin.

### Quantitative Reverse Transcriptase Polymerase Chain Reaction

Injured spinal cords 500μm rostral and 300μm caudal to the lesion from 7-10 control or miR-200a inhibitor electroporated animals were micro dissected and pooled for each biological replicate. Total RNA was isolated using Trizol (Invitrogen) according to the manufacturer’s instructions. Subsequent cDNA was synthesized from 1μg of DNaseI (NEB) treated RNA using either High Capacity cDNA Reverse Transcription kit (Applied Biosystems) or miScript II RT kit (Qiagen). The qRT-PCR was carried out using Light Cycler 480 SYBR Green I Master (Roche). MicroRNA qRT-PCR was carried out with custom designed primers to conserved miRNAs (Qiagen) and custom primers from IDT were used to quantify axolotl mRNAs:

*18S*_F: CGGCTTAATTTGACTCAACACG

*18S*_R: TTAGCATGCCAGAGTCTCGTTC

*brachyury*_F: GAAGTATGTCAACGGGGAAT

*brachyury*_R: TTGTTGGTGAGCTTGACTTT

*sox2*_F: TTGTGCAAAATGTGTTTCCA

*sox2*_R: CATGTTGCTTCGCTTTAGAA

*wnt3a*_F: AAGACATGCTGGTGGTCTCA

*wnt3a*_R: CCCGTACGCATTCTTGACAG

*wnt5a*_F: ACCCTGTTCAAATCCCGGAG

*wnt5a*_R: GGTCTTTGCCCCTTCTCCAA

*wnt8a*_F: TTGCTGTCAAATCAACCATG

*wnt8a*_R: TGCCTATATCCCTGAACTCT

*ctnnb1*_F: ACCTTACAGATCAAAGCCAG

*ctnnb1*_R: GGACAAGTGTTCCAAGAAGA

*lef1*_F: GTCCCACAACTCCTACCACA

*lef1*_R: TAGGGGTCGCTGTTCACATT

*fgf8*_F: TTTGTCCTCTGCATGCAAGC

*fgf8*_R: GTCTCGGCTCCTTTAATGCG

*fgf10*_F: AAACTGAAGGAGCGGATGGA

*fgf10*_R: TCGATCTGCATGGGAAGGAA

### Brachyury and Sox2 Probe Synthesis

Approximately 500bp fragments of axolotl *brachyury* and *sox2* were PCR amplified using OneStep PCR Kit (Qiagen) from RNA extracted from axolotl embryos at various developmental stages using the following primers:

*brachyury* ISH For: CCCCAACGCCATGTACTCTT

*brachyury* ISH Rev: GGCCAAGCGATATAGGTGCT

*sox2* ISH For: TGGCAATCAGGAAGAAAGTC

*sox2* ISH Rev: GCAAATGACAGAGCCGAACT

The resulting PCR fragments were gel purified using the Monarch Gel Purification Kit (New England Biolabs) and TA cloned into pGEM-T Easy (Promega) then transformed into DH5α competent *E. coli* (Invitrogen). Blue/White positive selection was used to pick clones and recovered plasmids were sent for sequencing. Positive clones were digested with the appropriate enzyme to linearize the plasmid and anti-sense ribonucleoprobe synthesis was carried out using Sp6 or T7 RNA polymerase (New England Biolabs)+DIG labeled UTP (Roche). Subsequent probes were cleaned up using the RNA Clean Up kit (Qiagen) and resuspended in hybridization buffer.

### Fluorescent In situ Hybridization

All RNAscope® *in situ* hybridization procedures were performed according to the manufacturer’s instructions (Advanced Cell Diagnostics). In brief, cryosections were incubated in PBS for 10 minutes to remove the OCT, and then baked at 60°C for 30 minutes. The slides were next post-fixed in 4% paraformaldehyde for 15 minutes at 4°C, and then dehydrated in a graded series of ethanol dilutions before being incubated in absolute ethanol for 5 minutes. After briefly air-drying the slides for 5 minutes, sections were next treated with hydrogen peroxide to quench endogenous peroxidase activity for 10 minutes at room temperature. Next, samples were briefly washed in deionized water, then incubated in target retrieval buffer at 90°C for 5 minutes. Following target retrieval, the slides were rinsed in deionized water for 15 seconds and treated with absolute ethanol for 3 minutes. Slides were next permeabilized in protease III for 30 minutes before hybridization with RNAscope® probes at 40°C for 2 hours. Following hybridization, sections were placed in 5x SSC overnight. The next day, sections were incubated in Amp1 and Amp2 at 40°C for 30 minutes each, followed by Amp3 for 15 minutes. Next, slides were treated with HRP-C1 to detect *brachyury* or *fgf10*, followed by a 30-minute incubation in OpaI-690 fluorescent dye. After treatment with HRP blocking buffer, samples were next incubated in HRP-C2 to detect either *sox2* or *wnt5a*, followed by a 30-minute incubation in OpaI-570 dye. After an additional treatment with HRP blocking buffer, slides were counterstained with DAPI and imaged using a Zeiss 780 Confocal Microscope.

### Lineage Tracing

Cells of the uninjured spinal cord were transfected with a construct containing a GFP or tdTomato fluorescent protein under the control of the axolotl GFAP promoter. The cells were injected and electroporated as previously described (Echeverri and Tanaka, 2003a, Sabin et al., 2015a). One day after electroporation the animals were screened for fluorescent cells. Positive animals were then injected with a control inhibitor or miR-200a inhibitor and then a spinal cord lesion performed as described in (Sabin et al., 2019). Animals were imaged every 3 days until the lesion site was no longer visible and the animals regained motor and sensory function, typically 12-14 days post injury.

### Cloning 3’ Untranslated Regions for miRNA Luciferase Assays

For 3’ UTR luciferase experiments, primers were designed to amplify the *brachyury* and *β-catenin* 3’ UTR based off sequences obtained from axolotl-omics.org. All the 3’ UTRs were amplified with a 5’ SpeI and 3’ HindIII restriction site.

*brachyury* 3’ UTR For 1 AGCACTAGTATGTGAAATGAGACTTCTAC

*brachyury* 3’ UTR Rev 1 TGCAAGCTTCTTATTCTTCCCATTTAACTTAAA

*ctnnb1* 3’ UTR For 1 ATAACTAGTTTGTGTAATTTTTCTTAGCTGTCATAT

*ctnnb1* 3’ UTR Rev 1 ATCAAGCTTAATTGCTTTATAGTCTCTGCAGAT

*ctnnb1* 3’ UTR SDM1 For AGTGCCTGATGAATTCAACCAAGCTGAG

*ctnnb1* 3’ UTR SDM1 Rev CTCAGCTTGGTTGAATTCATCAGGCACT

*ctnnb1* 3’ UTR SDM2 For ATTTAATGGTGTAGGAATTCAATAGTATAA

*ctnnb1* 3’ UTR SDM2 Rev TTATACTATTGAATTCCTACACCATTAAAT

The PCR fragments and pMiR Report (Life Technologies) were digested with SpeI and HindIII (NEB) and the fragments were ligated over night at 4°C with T4 DNA Ligase (NEB) and heat shock transformed into DH5α competent *E. coli* (Promega).

### Mutation of miR-200 sites in Brachyury 3’ UTR

To mutate the 3 miR-200a and 3 miR-200b sites in the axolotl Brachyury 3’ UTR, we used the QuikChange Lightening Multi Site-Directed Mutagenesis kit (Agilent) as per the manufacturer’s instructions.

miR-200a1 SDM: gactgctttctatggacactttttaatttctgaagataagctcccacccg

miR-200a2 SDM: cacacataaatcttttcgtgctgaacaaattatgatccatgaaaccagtgcatcattt

miR-200a3 SDM: tccaatgtgtgtaatcctctcaattatcgcctctgcgtgtagaatgtc

miR-200b1 SDM: atgcattacaatgcattgttttctggacggcaatgaaagctgtgatgaaatatttaagat

miR-200b2 SDM: caccataagagacaataaatgcaccggaatactgtgatatttgatgcctgcac

miR-200b3 SDM: gaatcattaccatgtatttatcaggccggaatattcaaaatgtgacttcctctgtga

### 3’ UTR luciferase experiments

B35 neuroblastoma cells were plated in a 96 well plate (Celltreat Scientific Products) at a concentration of 2.0 x 10^5^ cells/mL and allowed to adhere overnight. The next day cells were co-transfected with the appropriate Luciferase 3’ UTR reporter plasmid, β-Galactosidase control, and 100nM of miR-200a, miR-200b, or control mimic (Qiagen) per well using Lipofectamine 3000 (Invitrogen). After 48 hours luciferase activity was determined using Dual Light Luciferase & β-Galactosidase Reporter Gene Assay System (Thermo) according to the manufacturer’s protocol.

### Pie Chart and Venn Diagram Generation

Pie charts were generated using previously published data (Sabin et al., 2019) to represent the total number of differentially expressed genes in a given comparison using Excel. Venn Diagrams were generated with Venny (v2.1.0) (Oliveros 2007-2015) and saved as .csv files to be modified in Adobe Illustrator.

### Gene Ontology (GO) Analysis

GO terms were determined using GOrilla (Eden et al., 2009). We used 2 unranked list of genes: a background list (all differentially expressed genes in our data set) and a target list (genes that were differentially regulated in a given comparison). Using this approach, GOrilla generated a list of enriched Biological Process GO terms and we selected the top 9-13 terms with the lowest p-value and generated representative bar graphs using Excel.

### Calculation of the proportion and distribution of neural stem cells and newborn neurons

The number of Sox2^+^ neural stem cells were counted in control and miR-200a inhibitor spinal cords 2-weeks post injury. The proportion of Sox2^+^ neural stem cells was calculated by dividing the total number of Sox2^+^ neural stem cells by the total number of DAPI^+^ spinal cord cells times 100. To analyze regenerative neurogenesis, control or miR-200a inhibitor animals were injected with EdU at 5 and 7-days post injury, then tails were harvested for cryosectioning at 14 days post injury. The proportion of newborn neurons was determined by dividing the number of NeuN^+^/EdU^+^ double positive neurons by the total number of NeuN^+^ neurons times 100.

To visualize whether the proportion of Sox2^+^ neural stem cells changed rostral and caudal to the injury site in control and miR-200a inhibitor spinal cords, we quantified the average number of Sox2^+^ neural stem cells at a defined distance rostral and caudal from the lesion. We divided that number by the average number of DAPI^+^ spinal cord cells at that same distance and then binned the vales from 3 adjacent serial sections encompassing a region of 80μm. We used the same method for graphing the distribution of newborn neurons relative to the injury site.

### Statistical Analyses

All results are presented as mean +/- s.d. unless otherwise stated. Analyses were performed using Microsoft Excel or GraphPad Prism v8. Data set means were compared using ANOVA for three or more tests with a Tukey test (for multiple comparisons) or Dunnett test (to compare to a control mean). When two groups were compared an unpaired *t-*test was used. When multiple comparisons were made using a t-test, an adjusted p-value was determined using the two stage Benjamin, Krieger, and Yekutieli procedue with a false discovery rate <5%. Differences between groups was considered significant at three different levels (p-values of *≤0.05, **≤0.01 and ***≤0.001) and are indicated in the figure legends.

## Acknowledgements

We thank Ricardo Zayas for feedback on the manuscript.

## Data Availability

The authors declare that all data supporting the findings of this study are available within the article and its Supplementary Information files or from the corresponding author upon reasonable request. The RNA seq data has been deposited in the public GEO database with the accession number GSE122939.

**Supplementary Figure 1**: miR-200a affects expression of common and unique gene sets in the uninjured and regenerating spinal cord. (A, B, C) Pie chart representation of the proportion of up-regulated (Red) or down-regulated (Blue) genes in (B) uninjured control compared to uninjured miR-200a inhibitor electroporated spinal cords or (A) 4 days post injury control compared to 4 days post injury miR-200a inhibitor electroporated spinal cords. (C). Pie chart illustrating the number of overlapping versus individual genes that are differentially regulated. (B-F) Gene Ontology terms enriched in gene specifically in (B) control regenerating or (E) genes common to all data sets or specific to uninjured tissue (F).

**Supplementary Figure 2**: Multiple miR-200 members directly regulate the *brachyury* 3’ UTR. (A) Co-transfection of B35 cells with a *brachyury* 3’ luciferase reporter and a miR-200a (A) or miR-200b (B) mimic results in decreased luciferase activity compared to the control mimic (n=5). Mutation of all miR-200a seed sequences in the *brachyury* 3’ UTR alleviates this repression suggesting it is a direct target of miR-200 in axolotl (n=4). *** p≤0.001, N.S. is not significant. Error bars represent ±S.T.D.

**Supplementary Figure 3**: miR-200a may regulate the expression of Wnt signaling components. (A) qRT-PCR analysis revealed that miR-200a inhibition leads to increased expression of Wnt signaling transcriptional components *lef1* and *β-catenin*. (B) Co-transfection of B35 cells with a *β-catenin* 3’ UTR luciferase reporter and a miR-200a mimic leads to decreased luciferase activity compared to controls. Mutation of both miR-200a seed sequences in the *β-catenin* 3’ UTR alleviates this repression (n=3). *p≤0.05, *** p≤0.001, N.S. is not significant. Error bars represent ±S.T.D.

**Supplementary Figure 4**: *brachyury* is expressed in regenerating spinal cord stem cells following amputation in larval animals. At 4 days post amputation (dpa) both *sox2* (ii) and *brachyury* (iii) are abundant in the spinal cord stem cells and share an overlapping expression pattern (iv)(n=6). Scale bar= 50mm.

