## Supplementary figures and images for "Regulation of stem cell identity by miR-200a during spinal cord regeneration"

### Supplemental Figures

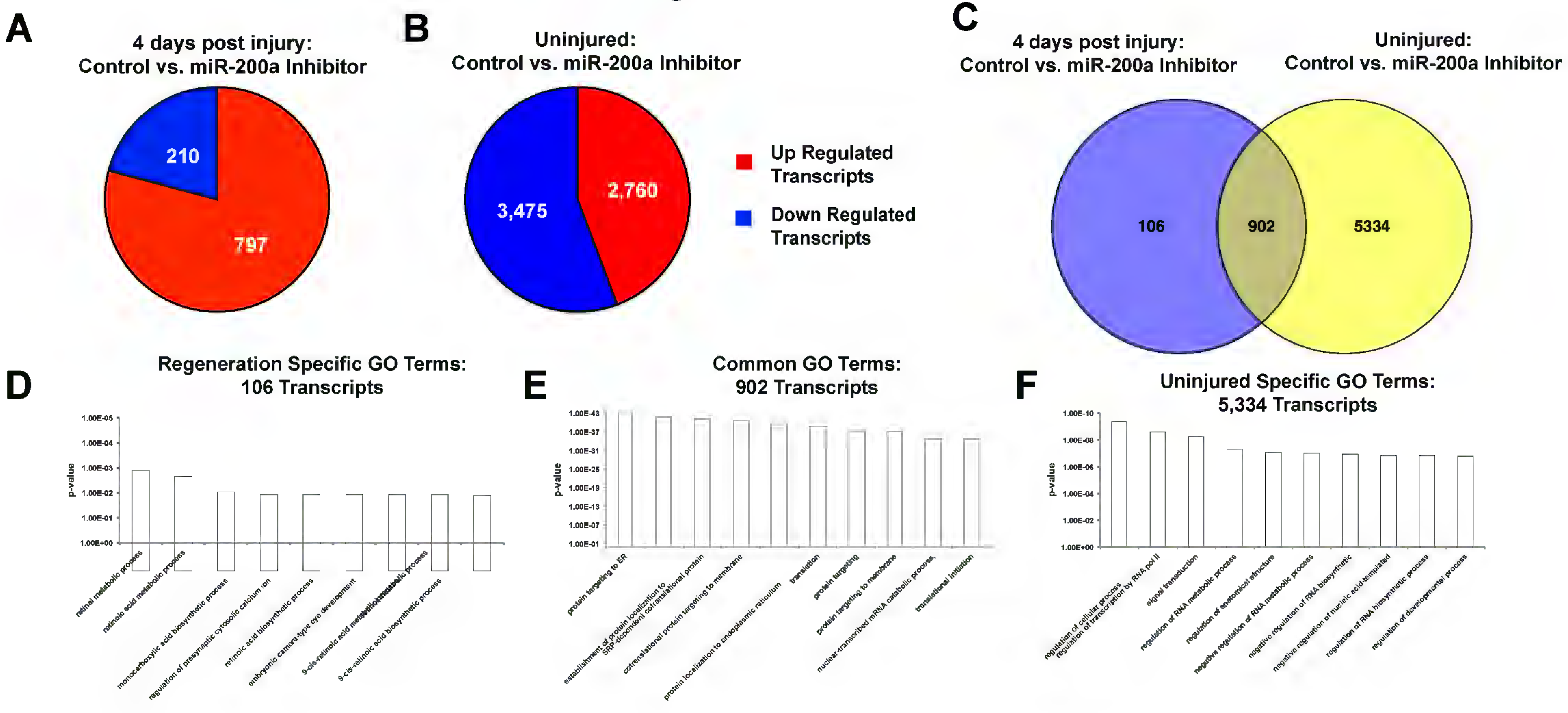

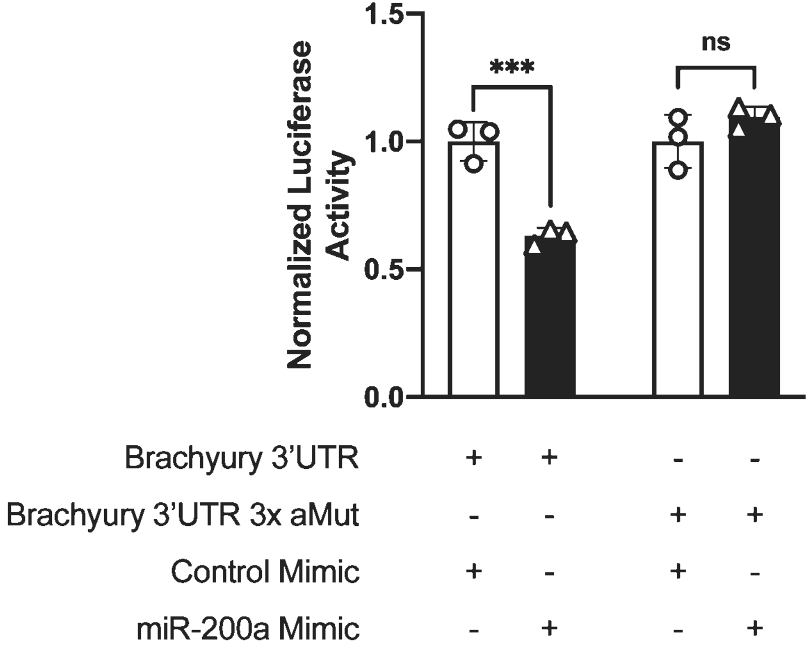

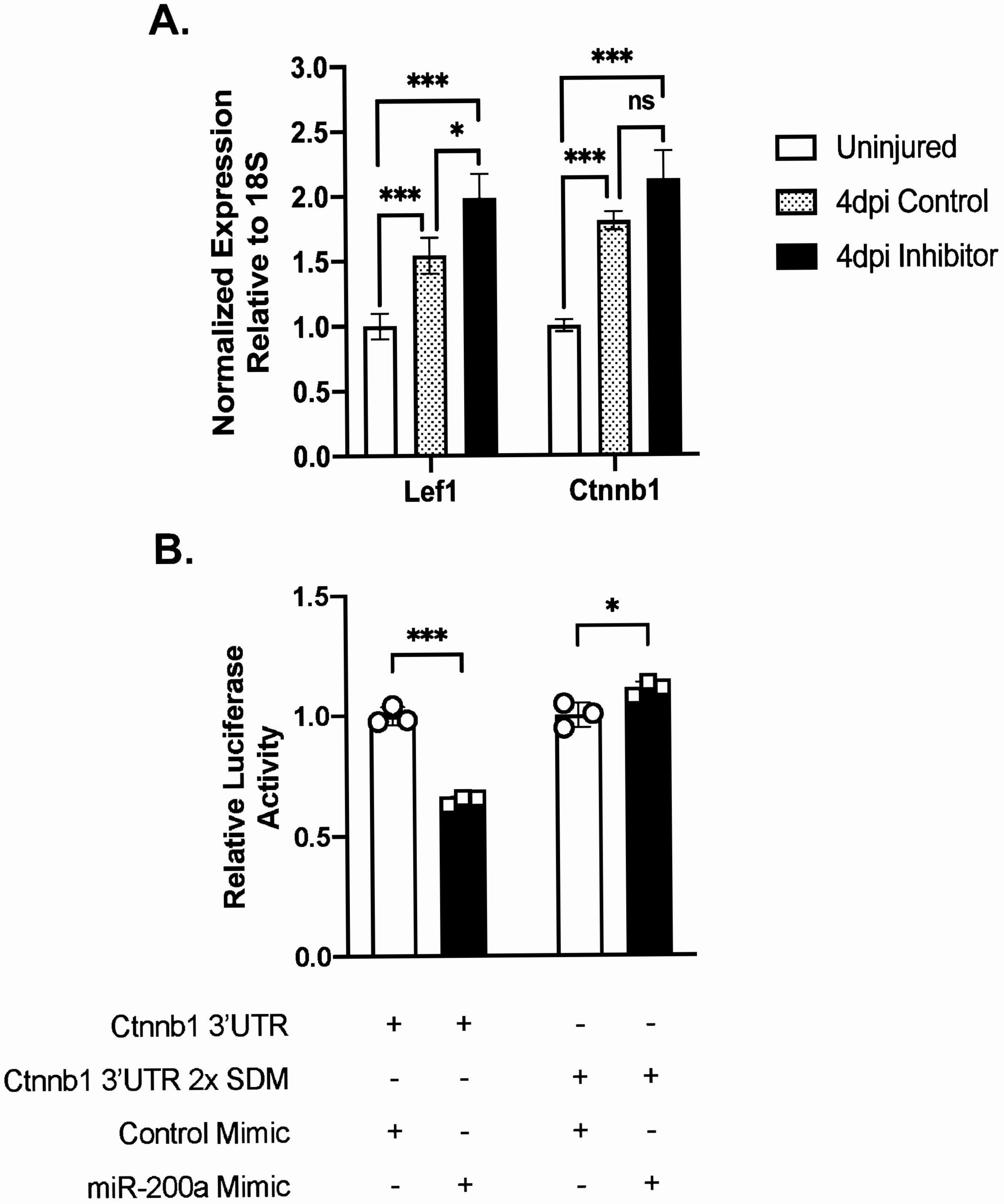

Figure S4

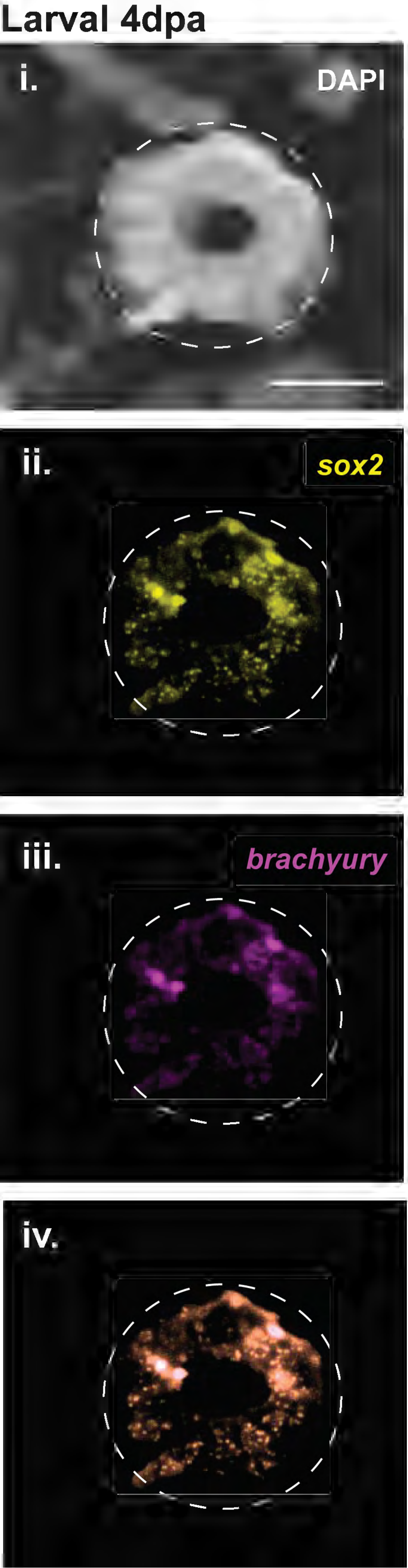
